# *VHL* Synthetic Lethality Signatures Uncovered by Genotype-specific CRISPR-Cas9 Screens

**DOI:** 10.1101/588707

**Authors:** Ning Sun, Sakina Petiwala, Charles Lu, Jessica E Hutti, Min Hu, Mufeng Hu, Marc H Domanus, Diya Mitra, Sadiya N Addo, Christopher P Miller, Namjin Chung

## Abstract

**Background:** Genome-wide CRISPR-Cas9 essentiality screening represents a powerful approach to identify genetic vulnerabilities in cancer cells. Here, we applied this technology and designed a strategy to identify target genes that are synthetic lethal (SL) with *von Hippel-Lindau* (*VHL*) tumor suppressor gene. Inactivation of *VHL* has been frequently found in clear cell renal cell carcinoma (ccRCC). Its SL partners serve as potential drug targets for the development of targeted cancer therapies.

**Results:** We performed parallel genome-wide CRISPR screens in two pairs of isogenic ccRCC cell lines that differ only in the *VHL* status. Comparative analyses of screening results not only confirmed a well-known role for mTOR signaling in renal carcinoma, but also identified DNA damage response and selenocysteine biosynthesis pathways as major SL targets in *VHL-*inactivated cancer cells. Follow-up studies provided cellular and mechanistic insights into SL interactions of these pathway genes with the *VHL* gene.

**Conclusions:** Using isogenic CRISPR screening approach, we uncovered novel biological processes that are SL with *VHL*, which can be exploited for drug development for ccRCC. Our CRISPR and RNA-seq datasets provide a rich resource for future investigation of the function of the VHL tumor suppressor protein. Our work demonstrates the efficiency of CRISPR-based synthetic lethality screening in human isogenic cell pairs. Similar strategies could be employed to unveil SL partners with other oncogenic drivers.

## Background

Kidney cancer is one of the most common cancers in both men and women, with ccRCC being the most frequent and aggressive subtype [1, 2]. Biallelic inactivation of the *VHL* gene caused by chromosome 3p loss, mutation or hypermethylation is observed in over 80% of ccRCC [3, 4]. The VHL protein, pVHL, functions as the recognition subunit of an E3 ubiquitin ligase complex. The canonical role of pVHL is to promote the degradation of the hypoxia-inducible transcription factors α (HIF1α, HIF2α, and HIF3α, collectively HIFα) through oxygen-sensitive ubiquitin-mediated proteolysis. Loss of function of pVHL results in the constitutive stabilization of HIFα, which promotes inappropriate expression of downstream genes involved in angiogenesis, metabolism, and proliferation and thus contributes to tumorigenesis [5]. In addition to its role in HIFα regulation, pVHL has been implicated in a variety of HIF-independent processes including extracellular matrix regulation [6], cilium formation [7–10], microtubule stabilization [11], protein synthesis [12], apoptosis [13], mitosis [14, 15], cell senescence [16, 17] and DNA repair [18]. With increased understanding of pVHL function and ccRCC pathogenesis, a number of targeted therapies have been developed, including antiangiogenic agents and mTOR inhibitors [19]. However, the development of drug-induced resistance and dismal prognosis of patients with metastatic diseases has heightened the need for novel therapeutics for ccRCC management.

Since inactivation of *VHL* is a signature lesion in almost all ccRCC, it provides the opportunity to discover *VHL-*specific genetic vulnerabilities in tumor cells that can be exploited for discovery of new therapeutic targets. Since it is challenging to pharmacologically restore an inactivated tumor suppressor gene, one approach would be to identify genes that are synthetically lethal (SL) with *VHL*. Two genes are considered to be SL if mutations of both genes lead to cellular or organismal death but the mutation of either gene alone enables viability [20, 21]. Therapeutic strategies based on synthetic lethality may provide drug targets when the cancer-causing genes cannot readily be targeted and enable selective killing of cancer cells while sparing normal cells [22, 23]. To this end, several *VHL* SL partners have been discovered using multiple approaches including hypothesis-driven mechanistic studies, compound screening and small-scale short hairpin RNA screening [24–36]. However, systematic identification of *VHL* SL genes at a genome-wide scale has not been reported to date.

Clustered regularly interspaced short palindromic repeat (CRISPR)/CRISPR-associated protein 9 (Cas9) has recently emerged as a powerful and versatile tool for mammalian genome engineering [37–42]. The site-specific DNA double-strand breaks induced by CRISPR-Cas9 are mainly repaired by non-homologous end joining, which often causes gene disruption by frameshift mutation. Given its high efficiency and scalability, CRISPR-Cas9 has been quickly adopted in genome-wide loss-of-function genetic screens to identify common and context-dependent essential genes in mammalian cells [43–53]. Here, we developed a strategy to identify *VHL* SL genes by performing parallel CRISPR-Cas9 screens in isogenic pairs of cancer cell lines that are genetically identical except for the *VHL* status. Through genome-wide CRISPR screening followed by rigorous validation screening and transcriptome profiling, we identified novel biological processes that are SL with *VHL*, including selenocysteine biosynthesis and DNA damage response pathway. We validated screening hits using genetic and pharmacological approaches and elucidated underlying molecular mechanisms. We also discovered a subset of the *VHL* SL genes had elevated expression in *VHL-*deficient cell lines and in primary ccRCC tumor tissues, which correlated with poor overall survival of affected patients. Our work uncovers paths to novel therapeutic strategies for ccRCC and provides a rich resource for future investigation of the function of the VHL tumor suppressor protein. Similar approaches could potentially be applied to identify SL partners of other oncogenic drivers such as PI3K pathway alterations, *MYC* activation, *TP53* inactivation and *RAS* oncogenes.

## Results

### Generation of isogenic *VHL* ccRCC cell line pairs

To facilitate the discovery of genetic vulnerabilities dependent on *VHL* loss, we took an unbiased genome-wide CRISPR screening approach using isogenic pairs of ccRCC cell lines that only differ in the *VHL* gene (Fig. 1A). To enable this approach, we transduced two widely-used *VHL-*deficient ccRCC cell lines, A-498 and 786-O, with lentiviral constructs expressing either pVHL co-translated with Cas9 or Cas9 alone (hereinafter referred to as A-498-*VHL*^+^ and 786-O-*VHL*^+^, and A-498-*VHL*^−/−^ and 786-O-*VHL*^−/−^, respectively; Fig. 1B). The engineered cell lines exhibited consistent and high levels of Cas9 activity in *EGFP* knockout (KO) reporter assays (Supplemental Fig. S1). We further validated these cell lines for functional restoration of the *VHL* gene by RNA-seq analysis (Supplemental Table S1,S2). Compared to *VHL*^−/−^ cells, *VHL*^+^ cells showed upregulation of the genes in Krebs cycle and p53 signaling pathway, and downregulation of those associated with hypoxia and NF-kB signaling (Supplemental Fig. S2). These observations are consistent with the established role of pVHL in both normal cells and ccRCC tumors [13, 54–58]. In agreement with previous findings [59–61], *VHL* status did not affect the rate of cell proliferation under normal cell culture conditions (Supplemental Fig. S3). Overall, we created two pairs of Cas9-proficient *VHL*^−/−^ and *VHL*^+^ isogenic cell lines, which recapitulated key aspects of known ccRCC biology and served as cellular models for *VHL* SL CRISPR screens.

**Figure 1.**
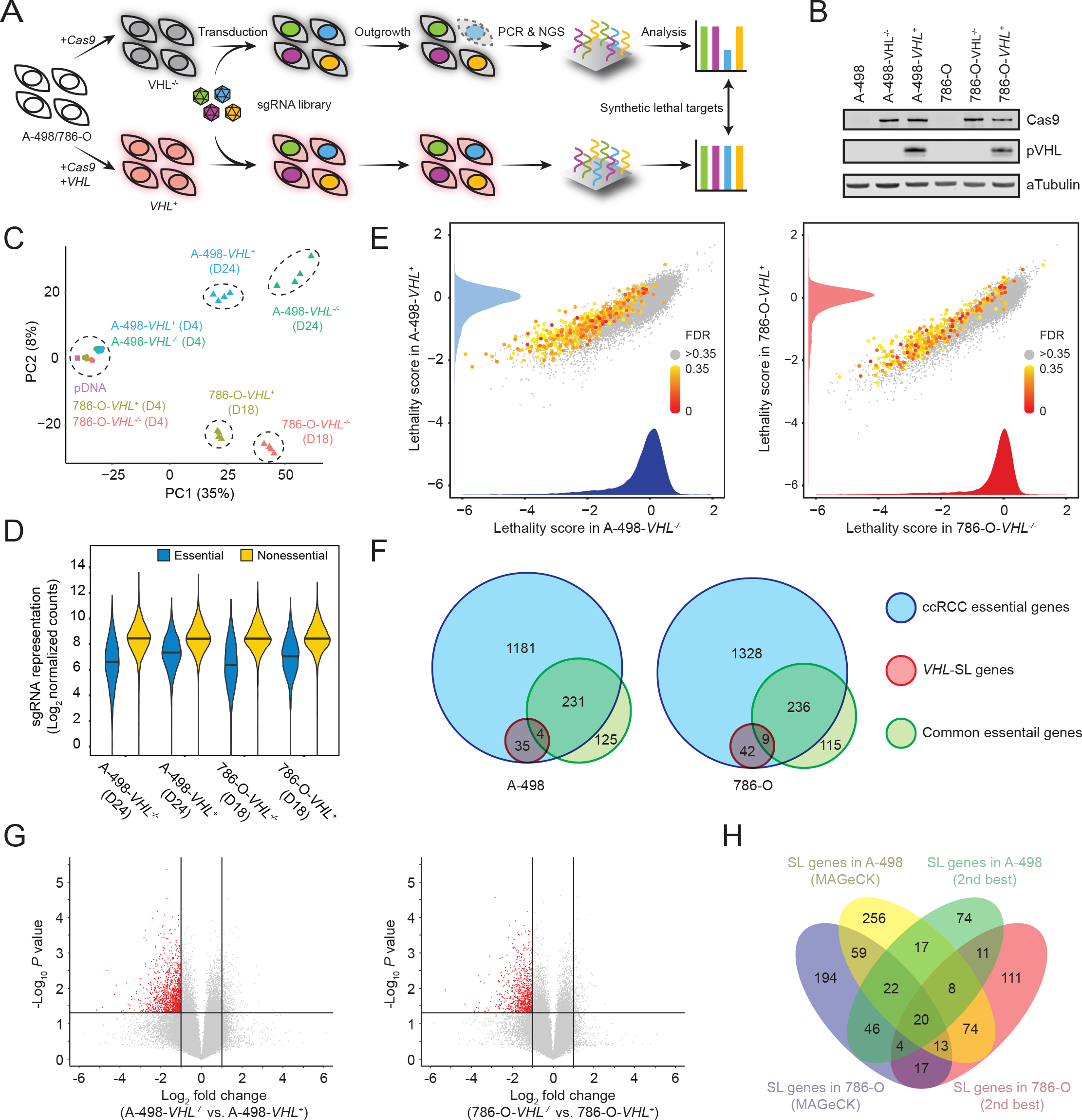
Genome-wide CRISPR-Cas9 synthetic lethality screens in isogenic ccRCC cell line pairs. (A) Workflow and screening strategy of CRISPR-Cas9 synthetic lethality screening. (B) Western blot confirmation of Cas9 and pVHL expression in designated cell lines. (C) Principal component analysis of sgRNA-sequencing results of biological replicates at various time points. (D) Violin plot of the representation of sgRNAs targeting a reference set of essential and nonessential genes [63]. (E) Gene lethality scores of all genes screened in two pairs of isogenic ccRCC cell lines. Each gene is colored according to its MAGeCK FDR based on synthetic lethality comparing *VHL*^−/−^ to *VHL*^+^ cells. Inset graphs show lethality score distributions. (F) Venn diagram of ccRCC cell line-specific essential genes, *VHL* SL genes and common essential genes [63]. (G) Volcano plots of significance (*P* value) versus fold change for differential sgRNA depletion analysis. (H) Venn diagram of *VHL* SL gene lists isolated from two isogenic cell line pairs using two different hit selection strategies.

### Genome-wide CRISPR-Cas9 screens in isogenic *VHL* ccRCC cell lines

We subsequently proceeded to perform CRISPR screens in the two pairs of isogenic ccRCC cell lines using a genome-wide lentiviral library [62]. After large-scale lentivirus transduction, cells were passaged serially under optimal conditions and a statistically sufficient number of cells were sampled at an early passage (4 days post-transduction) and a late passage (24 or 18 days post-transduction for A-498 and 786-O, respectively). sgRNAs were deconvoluted and quantified through deep sequencing. Screening results demonstrated high reproducibility and strong correlations among four replicates (Pearson’s r=0.95 and 0.80 for the early and late passages, respectively; Supplemental Fig. S4,S5). Principal component analysis (PCA) of screening results showed the largest variance along population doubling number or the duration in cell culture, with all early-passage cells clustering together with each other and with plasmid DNA library (pDNA) on one end, and late-passage cells forming distant, separate clusters on the other end (PC1, 35%; Fig. 1C). Different cellular makeups between A-498 and 786-O manifested during late passages, forming the second principal component (PC2, 8%). As expected, the sgRNAs targeting a curated set of common essential genes were significantly depleted during late passages, compared to non-targeting control sgRNAs or those targeting nonessential genes [63] (Fig. 1D and Supplemental Fig. S6).

For each gene in each cell line, we defined its lethality score as the median of log2 fold change in the abundance of all sgRNAs targeting the same gene comparing late passage to pDNA and calculated its false discovery rate (FDR) using MAGeCK algorithm (Fig. 1E and Supplemental Table S3) [64]. Using FDR<0.1 as the cutoff, 1,451 and 1,615 genes were identified as essential genes for cell fitness in A-498 and 786-O cell lines, respectively. We then compared lethality scores between *VHL*^−/−^ and *VHL*^+^ cells to identify *VHL* SL genes. When an FDR<0.1 cutoff was applied, only 39 genes in A-498 and 51 genes in 786-O deemed *VHL* SL, and they were largely different from the curated set of common essential genes (Fig. 1F). To validate screening hits and minimize false negative discovery, we applied a more relaxed FDR of 0.35, and identified 469 and 375 genes as A-498 and 786-O *VHL* SL candidate genes, respectively (Fig. 1E). We supplemented our MAGeCK-based hit selection with a second, independent method that evaluated the “second-best” sgRNA of each gene for selective lethality for *VHL*^−/−^ cells over *VHL*^+^ cells using log2 fold change <-1 and *p*<0.05 as a cutoff (Fig. 1G and Supplemental Table S4). The “second-best” method identified 258 and 202 genes for A-498 and 786-O cells, respectively. Combining these genes together, we generated a total of 926 genes for further validation (Fig. 1H and Supplemental Table S5).

### Validation of screening hits with orthogonally designed sgRNAs and gene expression profiling

A new sgRNA library was constructed for validation of the 926 candidate *VHL* SL genes by selecting ten new orthogonal sgRNAs per gene, based on an improved sgRNA picking algorithm (11,190 sgRNAs targeting the 926 primary screening hits as well as 1,890 negative control sgRNAs) [62]. Validation screens were performed in a fashion similar to primary screens, except that intermediate-passage samples were also collected (Day 4 for early passages; Day 14 or 11 for intermediate-passage A-498 or 786-O cell lines, respectively; and Day 23 or 18 for late-passage A-498 or 786-O cell lines, respectively). Results of the PCA analysis were similar to those obtained in the primary screen, with an additional granularity introduced by intermediate-passage samples (Fig. 2A). Further, the majority of the genes in validation screens showed significantly reduced viability for *VHL*^−/−^ compared to *VHL*^+^ cells at intermediate and late passages, suggesting these genes exhibited a SL phenotype (Fig. 2B). Synthetic lethality scores were defined as the median of log2 fold change in the abundance of all sgRNAs targeting the same gene between *VHL*^−/−^ and *VHL*^+^ cells at late passages. Despite relaxed hit selection approaches involving moderately high FDR (0.35) and selection of hits by two different methods, the synthetic lethality scores showed high concordance between primary and validation screens (Fig. 2C and Supplemental Table S6). In order to remove as many false positives as possible, a more stringent MAGeCK FDR of 0.05 was applied to the validation screening data. This slightly reduced gene counts from 612 to 582 for A-498 cell lines, and from 485 to 384 for 786-O cell lines (Fig. 2D and Supplemental Table S7). Potential false positives were further eliminated by removing the genes that did not appear to be expressed in these cell lines based on RNA-seq data. By applying fragments per kilobase of transcript per million mapped reads (FPKM) of less than 0.1 as a cutoff, 36 (6%) and 14 (4%) genes for A-498 and 786-O cell lines, respectively, were eliminated (Supplemental Table S8). Together, these strategies allowed us to define a set of 350 *VHL* SL genes that were both expressed and functional in A-498 and 786-O cell lines (Fig. 2D and Supplemental Table S9).

**Figure 2.**
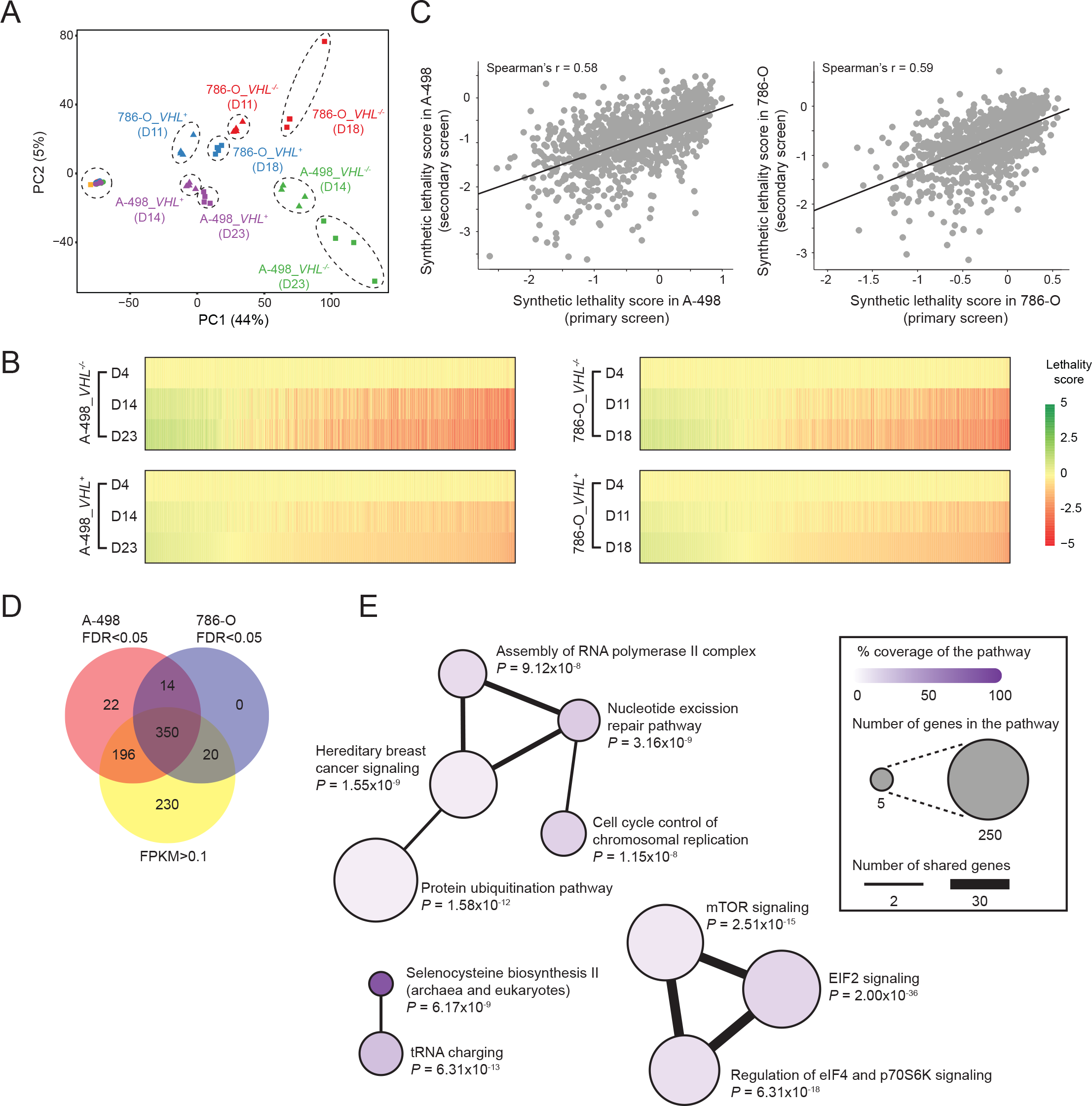
Secondary CRISPR screens revealed novel *VHL* SL biological pathways. (A) Principal component analysis of sgRNA-sequencing results of biological replicates at various time points for secondary CRISPR screening. (B) Heatmaps of gene lethality scores across two pairs of ccRCC cell lines at the indicated time points. (C) Scatter plots of gene synthetic lethality scores in primary and secondary CRISPR screens. (D) Hit triage using secondary CRISPR screening and RNA-seq data sets. (E) Most significant pathways by Ingenuity Pathway Analysis from the 350 *VHL* SL genes. *P* values were derived from Fisher’s exact tests.

Gene ontology and pathway analyses of these 350 genes confirmed a previously known interaction between *VHL* and mTOR signaling (Fig. 2E and Supplemental Table S10). mTOR-related pathways are frequently dysregulated in ccRCC, mTOR inhibitors show clinical activities in advanced RCC patients, and *MTOR* is synthetically lethal with *VHL* [31] [65] [66, 67]. Furthermore, our analysis revealed two biological processes previously unknown to interact with *VHL*: selenocysteine biosynthesis and DNA damage response signaling. Given their high enrichment scores, we focused our attention on these two new pathways for target validation and further mechanistic investigations.

### Inhibition of selenocysteine biosynthesis is synthetically lethal with *VHL* loss

Selenocysteine, also known as the 21st amino acid, is encoded by a UGA codon and incorporated into selenoproteins using a unique tRNA (Fig. 3A) [68, 69]. Validation screening results demonstrated that sgRNAs targeting 5 of 6 genes (83%) in the selenocysteine biosynthetic pathway were selectively depleted in *VHL-*defective ccRCC cells compared to their *VHL-*proficient counterparts (Fig. 3B). We confirmed these observations with lentiviruses expressing individual sgRNAs. Knockout of the *SEPHS2* or *PSTK* using two independent sgRNAs led to selective inhibition of *VHL*^−/−^ cell proliferation (Supplemental Fig. S7). In contrast, cells transduced with scrambled, non-targeting control (NTC) sgRNAs did not exhibit meaningful differences between *VHL*^−/−^ and *VHL*^+^ cells. We went on to use a fluorescence-based competitive growth assay for more quantitative analysis of synthetic lethality (Fig. 3C). Briefly, we mixed violet-excited GFP (vexGFP)-positive cells expressing an NTC sgRNA with mCherry-positive cells expressing a gene-targeting sgRNA in equal ratio and allowed them to compete for the same pool of resources during an extended period of co-culture. If the gene-targeting sgRNA inhibited cell proliferation, then mCherry-positive cells would be depleted over time. Indeed, inactivation of the *SEPHS2* or *PSTK* gene using two different sgRNAs preferentially inhibited *VHL*-defective cells compared to *VHL*-proficient cells in both ccRCC cell line backgrounds (Fig. 3D).

**Figure 3.**
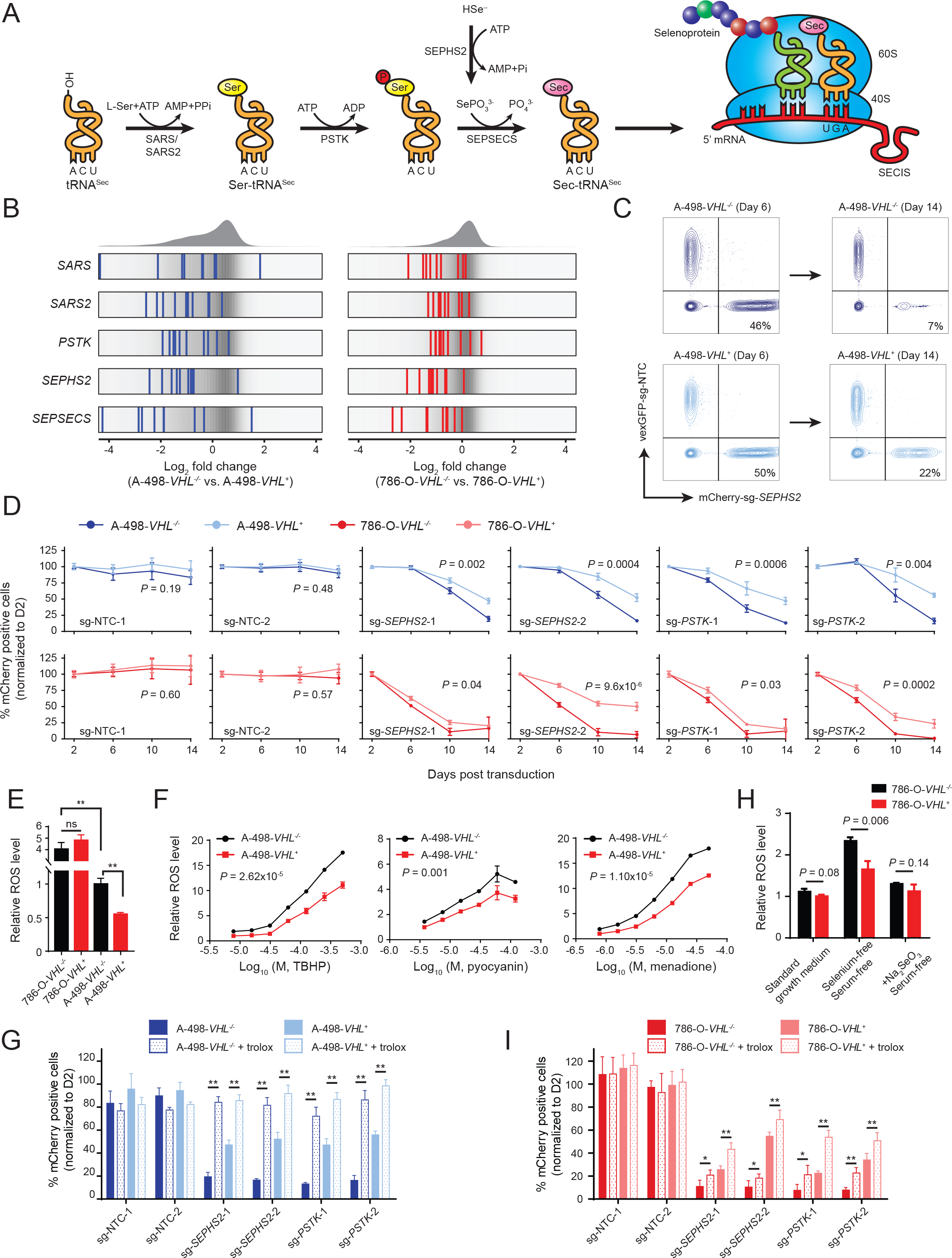
Inhibition of selenocysteine biosynthesis is synthetically lethal with *VHL loss*. (A) Biosynthesis of selenocysteine. The pathway begins with the attachment of serine to tRNA^Sec^ by seryl-tRNA synthetase (SARS) in cytoplasm or seryl-tRNA synthetase 2 (SARS2) in mitochondria. Phosphoseryl-tRNA kinase (PSTK) phosphorylates the serine moiety to form an intermediate, which in turn is converted to Sec-tRNA^Sec^ by selenocysteinyl-tRNA^Sec^ synthase (SEPSECS) using selenophosphate. Selenophosphate is synthesized by selenophosphate synthetase 2 (SEPHS2) using selenide. Finally, Sec-tRNA^Sec^ is delivered to the ribosome and inserted into selenoprotein. SECIS, an *in-cis* element in the selenoprotein mRNA located in the 3′-untranslated region, is required for decoding of the selenocysteine UGA codon. (B) Frequency histograms of the changes of sgRNA abundance in *VHL*^−/−^ versus *VHL*^+^ cells in secondary screening. sgRNAs targeting indicated genes are shown by the blue or red lines for A-498 or 786-O, respectively. (C) Representative flow cytometry plots of fluorescence-based competitive growth assay for the quantification of *VHL* synthetic lethality. (D) Genetic validation of the findings of the screen using competitive growth assay. Data represent mean ± SD (*n* = 4). *P* values were derived from *t* tests based on area under the curve. (E) Basal ROS levels of the isogenic cell lines (normalized by the ROS signal in A-498-*VHL*^−/−^ cells) in standard growth medium. Data represent mean ± SD (*n* = 12). *P* values were derived from *t* tests: \*\**P* < 0.005; ns, non-significant. (F) Dose-dependent ROS induction by *tert*-butyl hydroperoxide (TBHP), pyocyanin or menadione. Data represent mean ± SD (*n* = 3). *P* value was derived from *t* test of area under the curve. (G) Competitive growth assay in A-498 showed that treatment of trolox completely rescued growth inhibition induced by inactivation of *SEPHS2* or *PSTK*. Flow cytometry analysis was performed after treatment with 100 µM trolox or without trolox for at least 10 days in the competitive growth assay. Data represent mean ± SD (*n* = 4). *P* values were derived from *t* tests: \*\**P* < 0.005. (H) Effect of selenium deficiency on the ROS induction in the isogenic pair of 786-O cells. Cells were cultured in serum-free medium with or without 6.25 nM sodium selenite for 48 hours and the intracellular ROS levels were measured and normalized by the ROS signals in the 786-O-*VHL*^+^ cells cultured for 48 hours in standard growth medium. Data represent mean ± SD (*n* = 3). *P* values were derived from *t* tests. (I) Competitive growth assay in 786-O showed that treatment of trolox partially rescued growth inhibition induced by inactivation of *SEPHS2* or *PSTK*. Flow cytometry analysis was performed after treatment with 100 µM trolox or without trolox for at least 10 days in the competitive growth assay. Data represent mean ± SD (*n* = 4). *P* values were derived from *t* tests: \**P* < 0.05; \*\**P* < 0.005.

The human selenoproteome consists of 25 selenoproteins that are expressed in various tissues and organs [70]. Although the role of selenoproteins in tumorigenesis remains elusive, certain seleno-enzymes such as glutathione peroxidases and thioredoxin reductases are known to regulate cellular redox and protect cells from reactive oxygen species (ROS) [71, 72]. As pVHL plays important roles in the cellular response to changing oxygen levels and redox potentials [17, 73, 74], it can be hypothesized that *VHL-*defective cells are sensitive to the inhibition of selenocysteine biosynthesis because *VHL-*defective cells have dysfunctional ROS homeostatic mechanisms and develop greater dependence on selenoproteins in coping with the imbalanced cellular ROS for survival. In agreement with this hypothesis, we found that A-498-*VHL*^−/−^ cells had higher ROS signal than A-498-*VHL*^+^ cells at the basal level. (Fig. 3E). Furthermore, A-498-*VHL*^+^ cells more efficiently metabolized ROS generated by various ROS inducers such as *tert*-butyl hydroperoxide (TBHP), menadione and pyocyanin (Fig. 3F). Selenoprotein deficiency may be ameliorated by exogenously supplied ROS scavengers like trolox, a synthetic cellular antioxidant. Consistent with our hypothesis, treatment of trolox completely rescued A-498 cells from both *VHL* and selenoprotein deficiencies (Fig. 3G and Supplemental Fig. S8). Interestingly, 786-O cells displayed significantly higher levels of ROS than A-498 cells, and *VHL* restoration did not reduce their basal ROS levels in standard cell culture media (Fig. 3E). To assess the degree of contribution by selenoproteins to ROS homeostasis in *VHL*^−/−^ cells, we measured cellular ROS levels in the absence of selenium, an essential mineral that can be supplied in organic form as sodium selenite (Na_2_SeO_3_) to serum-free media. While A-498 cells could not be evaluated as they died in serum-free media (data not shown), 786-O cells showed more than two fold increase in ROS levels upon selenium deprivation, which was reversed by *VHL* restoration (Fig. 3H). When selenium was added back to serum-free media, ROS was further decreased to a level that could not be distinguished between *VHL*^−/−^ and *VHL*^+^ cells. Trolox treatment rescued 786-O cells from *VHL* deficiency, but not from selenoprotein deficiency (Fig. 3I and Supplemental Fig. S9). Taken together, the above data demonstrate critical cooperation between pVHL and selenoproteins in controlling cellular ROS homeostasis, and loss of both functions resulted in cancer cell death.

### DNA damage response components are synthetic lethal with *VHL*

Pathway analysis identified a second major class of *VHL* SL genes from DNA damage response (DDR) signaling [75–77], including nucleotide excision repair pathway, mismatch repair, DNA double-strand break repair, and cell cycle control (Fig. 2E and Supplemental Table S10). As pVHL plays a role in DDR and its depletion leads to the genomic instability in ccRCC [14, 15, 18], we hypothesized that *VHL*^−/−^ cells are burdened with a higher load of DNA damage that increases their requirement of DDR pathways for survival. Using cell proliferation and fluorescence-based competition assays, we confirmed that KO of DDR pathway genes (*CDK1*, *CHEK1* or *TOP2A*) selectively reduced the proliferation of *VHL*^−/−^ cells compared to *VHL*^+^ cells (Fig. 4A and Supplemental Fig. S10). We further assessed the synthetic lethality between DDR genes and *VHL* using small molecule inhibitors against AURKA, AURKB, ATR, CDK1 and TOP2A. Consistent with our genetic screening and validation studies, these compounds preferentially inhibited viability of *VHL*^−/−^ cells compared to their *VHL-*restored counterparts (Supplemental Table S11). RNA-seq analysis showed that the screening hits associated with DDR signaling were highly upregulated in *VHL*^−/−^ versus *VHL*^+^ cells (Supplemental Fig. S11 and Fig. 4B,C). Taken together, our data suggest that pVHL loss induces DNA damage-related stress, thus rendering *VHL*^−/−^ cells more reliant on DDR components for survival than *VHL*^+^ cells.

**Figure 4.**
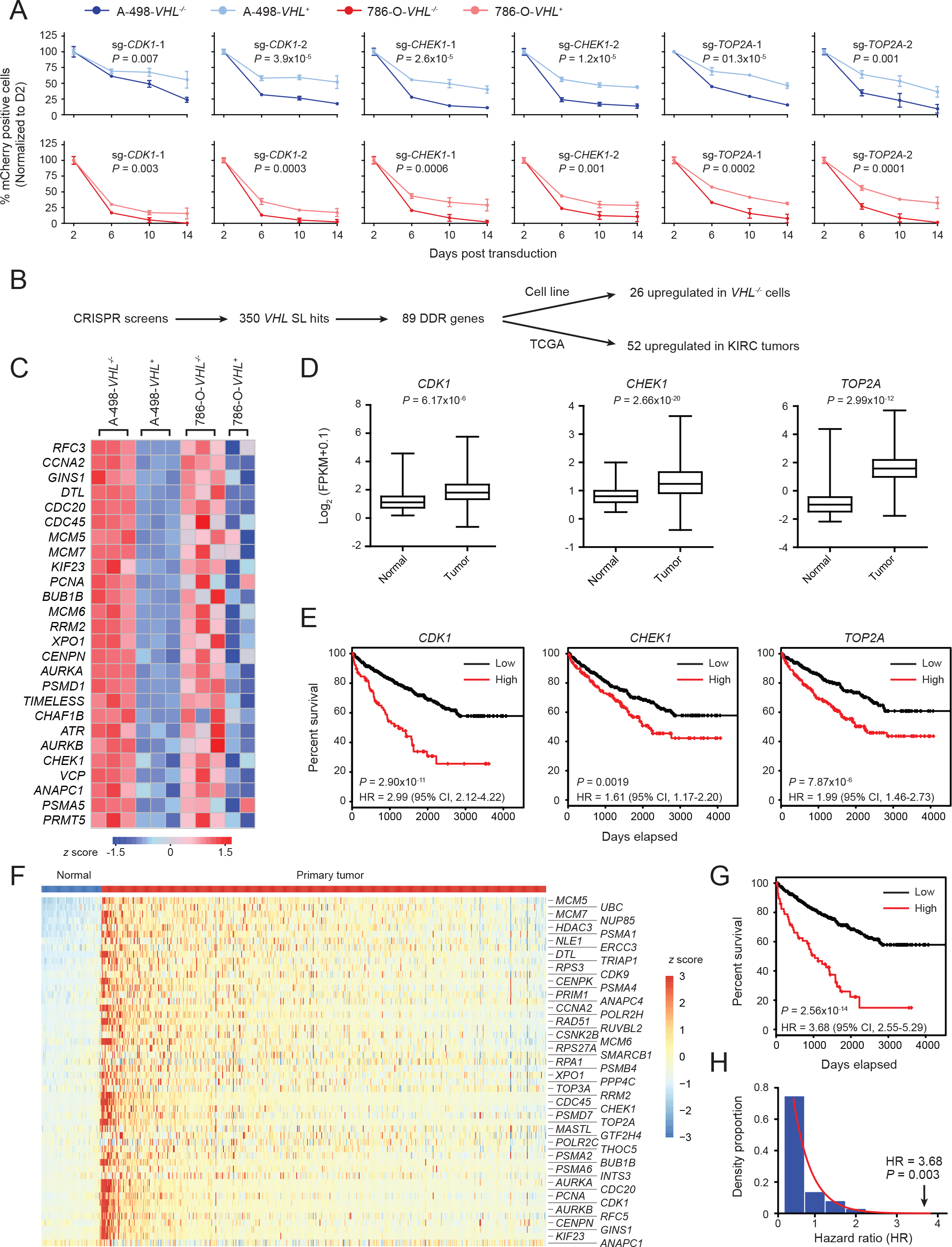
DNA damage response components are synthetic lethal with *VHL*. (A) Genetic validation of the selected DDR genes using competitive growth assay. Data represent mean ± SD (*n* = 4). *P* values were derived from *t* tests based on area under the curve. (B) Schematic for gene expression analysis of the *VHL* SL genes associated with DDR. (C) Heatmap of expression levels (*z* scores) of the upregulated *VHL* SL genes associated with DDR in isogenic ccRCC cell pairs. (D) mRNA expression of *CDK1*, *CHEK1* and *TOP2A* in primary tumors versus normal solid tissues from TCGA-KIRC cohort. *P* values were derived from *t* tests. (E) Survival analysis using the Kaplan-Meier method. Patients (TCGA-KIRC) were divided into two groups (high and low) based upon the mRNA expression levels, using the *z* score 1.96 as the cutoff. Hazard ratio (HR) and *P* values were derived from Cox’s proportional hazards analysis. (F) Heatmap of selected genes whose expression was significantly upregulated in primary tumors compared to normal tissues. (G) Survival analysis (Kaplan-Meier curve) comparing patients (TCGA-KIRC) with a high or low 52 gene signature score (using median *z* score 1.96 as the cutoff). HR and *P* values were derived from Cox’s proportional hazards analysis. (H) Distribution of HRs using randomly selected genes that are overexpressed in primary tumors versus normal tissues. TCGA gene expression data were applied with differential expression analysis. 1927 genes satisfied the criterion of selection (Log2 fold change >1, p<0.05). The non-overlapping groups, with each group composed of 52 distinct genes, were randomly selected to represent gene sets out of the 1927 genes. After being normalized into *z* scores, genes were selected according to each of the random gene set to perform survival analysis. This performance was repeated for 1000 times to generate 36000 HRs.

To explore the clinical relevance of our discovery, we analyzed The Cancer Genome Atlas Kidney Renal Clear Cell Carcinoma (TCGA-KIRC) data set, which includes transcriptome profiling of 537 primary tumors and 72 normal solid tissues correlated with clinical outcome of ccRCC patients [65]. Three of the validated screening hits, *CDK1*, *CHEK1* and *TOP2A*, were overexpressed in ccRCC (Fig. 4D) and associated with poor prognosis (Fig. 4E). Evaluation of all the DDR genes that we found to be SL with *VHL* showed that 52 DDR genes were significantly upregulated in primary ccRCC tumors (Fig. 4B,F and Supplemental Table S12). In addition, a gene expression signature derived from these 52 genes serves as a robust biomarker for predicting overall survival of ccRCC patients (Fig. 4G). In contrast, the gene expression signature of 52 randomly selected genes that are overexpressed in primary tumors was not associated with patient outcome (Fig. 4H). These results indicate that the screening hits we identified here represent *bona fide VHL* SL interactors in primary human tumors and serve as potential drug targets to develop effective treatment strategies.

## Discussion

Genome-wide *in vitro* cell proliferation screens based on CRISPR-Cas9 and RNAi approaches have been conducted in numerous cancer cell lines in order to identify and catalog genes essential for their survival, establishing several hundred “gold standard” genes commonly required for a number of cancer cell lines [50, 52, 53, 63, 78, 79]. Many such genes are also required for the survival of normal cells. As critical components of basic cellular metabolic and proliferative machinery, they provide limited insights into potential therapeutic strategy for cancer patients. One way to overcome such a limitation is to screen a panel of genetically related cancer cell lines and identify context-dependent essential genes that are required for some cancer cell lines carrying a specific mutation, but not others [51, 78, 79]. This approach often requires rather a large number of cancer cell lines, from tens to hundreds of them, to establish an unambiguous relationship between survival phenotype and a multitude of covariates such as genotypes, expression profiles, and tissue and cellular origins. Genotype-specific lethality or SL is a therapeutic strategy based on the observations that over the course of pathogenesis, cancer cells acquire a few key driver mutations and as a result become dependent on other genes for survival [80–82]. This strategy has been successfully employed in the development of PARP inhibitors as a novel therapeutic approach to certain cancers with BRCA mutations [83, 84]. *VHL* mutations present a promising opportunity to discover SL targets for ccRCC as they are found in up to 92% of the disease [4]. We hoped to reduce the number of cell lines needed to reveal SL genes by screening isogenic pairs of cell lines that are genetically identical except for a key tumor driver gene. To demonstrate this advantage, we utilized two *VHL-*null ccRCC cell lines and reconstituted them with a wild-type copy of the gene to create isogenic pairs of *VHL-* deficient and -proficient cell lines (Fig. 1A). Our screening data have shown that over one thousand genes are essential for the parental *VHL*-null ccRCC cell lines and they included more than two thirds of the “common essential” genes (Fig. 1F). However, the *VHL* SL genes we identified represented only a small fraction of genes essential for *VHL*-null ccRCC cell lines (2.7% and 3.2% for A-498 and 786-O cell lines, respectively) and furthermore differ from the “common essential” genes. These results demonstrate that use of isogenic cell line pairs allows for efficient identification of SL interactions, without the need to screen a large panel of heterogeneous cell lines.

ccRCC represents a serious unmet medical challenge as the prognosis for advanced patients remains poor. Better understanding of molecular genetic mechanisms that drive ccRCC could therefore lead to the development of efficient therapeutic strategies for this devastating disease. Our screens have uncovered multiple cellular signaling networks that are SL with *VHL*. Of these, mTOR signaling is one of the well-established therapeutic targets for ccRCC. However, two molecular networks, selenocysteine biosynthetic pathway and DDR pathway, have not previously been linked to this disease (Fig. 2E). Here, we show that five of the six selenocysteine biosynthetic pathway genes are SL with *VHL* (Fig. 3). Some selenoproteins are known to metabolize and neutralize ROS and are required for cellular defense mechanisms against oxidative stress. We observed that *VHL-*defective A-498 cells had significantly elevated levels of ROS compared to *VHL*^+^ isogenic cells under normal cell culture conditions and poorly metabolized ROS induced by exogenous agents (Fig. 3E,F). Therefore, both pVHL and selenoproteins play important roles in ROS homeostasis and maintaining cellular fitness in A-498 cells, which is consistent with the observation that a cell-permeable antioxidant, trolox, effectively rescued *VHL* inactivation and selenocysteine deficiency caused by *SEPHS2* or *PSTK1* KO. On the other hand, 786-O cells had constitutively high levels of ROS regardless of *VHL* status (Fig. 3E) but *VHL*^+^ isogenic cells better coped with ROS imbalance when selenium was depleted in cell culture (Fig. 3H). Consistent with these observations, overall survival of 786-O cells was more profoundly affected by selenocysteine deficiency but nevertheless *VHL*^+^ cells survived better than *VHL*^−/−^ isogenic cells (Fig. 3D), reflecting the greater competency of *VHL*^+^ cells to metabolize ROS in the absence of selenium (Fig. 3H). Trolox only partially rescued *VHL*^−/−^ and *VHL*^+^ 786-O cells (Fig. 3I), possibly because of more complex functions selenoproteins might have in the context of 786-O genetic makeup. It has been well appreciated that many types of cancer cell have increased levels of ROS. However, due to complex redox alterations and regulations, simply adding ROS-generating agents may not always lead to a selective killing of cancer cells [85]. Our discovery suggests that modulating unique redox regulatory mechanisms such as selenocysteine biosynthesis might provide a selective anticancer strategy.

Another major class of *VHL* SL genes belongs to DDR pathways (Fig. 4). Conventional chemotherapeutics like DNA alkylating agents kill cancer cells through mitotic catastrophe caused by cumulative DNA damage. However, use of these drugs has undesirable side effects because of a narrow therapeutic index. Our results show novel targets in DDR pathways with potentially wider therapeutic index. In addition to their potential as therapeutic targets, *VHL* SL DDR pathway genes showed strong predictive power for kidney cancer prognosis, especially when combined with mRNA profiling data (Fig. 4G). This suggests the importance of functional information provided by CRISPR screens in kidney cancer prognosis. Whereas the precise molecular basis of the SL interaction between *VHL* and selenocysteine biosynthesis or DDR will require further investigation, these studies suggest novel mechanisms of interplay between multiple cancer-relevant cellular processes. Interestingly, these two newly discovered *VHL* SL mechanisms may also be connected, as elevated ROS can cause DNA damage [85].

Although we focused on selenocysteine biosynthesis and DDR in this study, our screening results suggest additional *VHL* SL genes that merit further exploration. For example, protein ubiquitination pathway genes were significantly enriched among screening hits (Fig. 2E). As proteasome inhibitors are currently being used in clinic for cancer treatment [86], it would be interesting to investigate the efficacy of proteasome inhibitors in ccRCC. In another example, our screens identified a number of VHL *SL* genes associated with major metabolic pathways, including pyrimidine biosynthesis (*DTYMK* and *RRM2*), glycolysis (*TPI1* and *PKM*), fatty acid metabolism (*HSD17B10*), pentose phosphate pathway (*RPE*), and cholesterol biosynthesis (*GGPS1*). These observations reinforce already established roles of metabolic reprogramming in kidney cancers [87]. Finally, considering the increasing number of malignancies involving *VHL* inactivation, including hemangioblastoma, pheochromocytoma and pancreatic neuroendocrine tumors [88, 89], our study provides a framework for understanding molecular pathogenesis of *VHL* inactivation in additional cancer types.

## Conclusions

In this study, we applied a hypothesis-free strategy to identify *VHL* SL interactors based on CRISPR-mediated genetic screening. We performed parallel CRISPR screens in isogenic cell lines sharing the same genetic makeup except for *VHL* status. Through genome-wide screens followed by deeper focused screens, we confirmed a well-known role of mTOR signaling and uncovered novel biological processes, including selenocysteine biosynthesis and DNA damage response, which are SL with *VHL*. Our CRISPR and RNA-seq datasets provide a rich resource for future investigation of the function of pVHL. Our approach could potentially be employed to unveil SL partners with other oncogenic drivers.

## Abbreviations

VHL: von Hippel-Lindau
HIF: hypoxia-inducible transcription factor
ccRCC: clear cell renal cell carcinoma
SL: synthetic lethal
CRISPR: Clustered regularly interspaced short palindromic repeat
KO: knockout
PCA: Principal component analysis
pDNA: plasmid DNA
PC: principal component
FDR: false discovery rate
FPKM: fragments per kilobase of transcript per million mapped reads
ROS: reactive oxygen species
TBHP: tert-butyl hydroperoxide
DDR: DNA damage response
TCGA: The Cancer Genome Atlas

## Methods

### Cell culture

A-498 cells (ATCC, HTB-44) were cultured in Minimum Essential Medium Eagle (Sigma-Aldrich) plus 10% fetal bovine serum (FBS, ThermoFisher Scientific) and 100 U/mL penicillin/streptomycin (ThermoFisher Scientific). 786-O cells (ATCC, CRL-1932) were cultured in RPMI Medium 1640 (ThermoFisher Scientific) plus 10% FBS, 1 mM sodium pyruvate (ThermoFisher Scientific), 0.45% D-(+)-Glucose (Sigma-Aldrich) and 100 U/mL penicillin/streptomycin.

### Generation of Isogenic Cell lines for CRISPR Screening

786-O-*VHL*^−/−^ and A-498-*VHL*^−/−^ cells were generated by infecting parental cell lines with a lentiviral construct expressing *Cas9* and blasticidin-resistance gene in 12-well plates at 1000 xg for 2 hr, in the presence of 8 µg/µL polybrene (Sigma-Aldrich). Plates were then returned to 37 °C with 5% CO_2_. Cells were incubated overnight and then selected by blasticidin (5 μg/mL, ThermoFisher Scientific). 786-O-*VHL*^+^ and A-498-*VHL*^+^ cells were generated by infecting parental cell lines with a lentiviral construct expressing *Cas9*, *VHL* and blasticidin-resistance gene, followed by blasticidin selection (5 μg/mL).

### Western blotting

Cells were lysed directly in M-PER Mammalian Protein Extraction Reagent (ThermoFisher Scientific), separated on a NuPAGE 4-12% Tris-Glycine Gel (ThermoFisher Scientific), and transferred to a nitrocellulose membrane (Bio-rad). Membranes were blocked with Odyssey Blocking Buffer (LI-COR Biosciences) for one hour at room temperature. Membranes were then probed with anti-FLAG tag (1:2000, R&D Systems), anti-pVHL (1:500, Cell Signaling Technology), and anti-αTubulin (1:10,000, Novus Biologicals) primary antibodies at 4 °C. Membranes were washed four times in PBS-Tween (0.1%) before incubation in IRDye 680RD Goat anti-Mouse (1:10,000, LI-COR Biosciences) or IRDye 800CW Goat anti-Rabbit (1:10,000, LI-COR Biosciences) secondary antibody for one hour at room temperature. Membranes were washed four times in PBS-Tween (0.1%) before visualization using Odyssey CLx Imaging System (LI-COR Biosciences).

### RNA extraction and sequencing

Total RNA was extracted from 1-2 million cells using RNeasy Mini Kit (QIAGEN) using QIAcube (QIAGEN). RNA sequencing libraries were prepared with TruSeq Stranded mRNA library preparation kit (Illumina) according to manufacturer’s directions. Multiplex libraries were sequenced on a single read High Output NextSeq500 V2 flow cell (Illumina) with 150 cycles.

### RNA expression analysis

RNA-seq analysis was carried out by aligning Illumina reads using STAR aligner [90] against human reference sequence (Ensembl Release 84). Expression level is estimated using FeatureCount [91]. Differential Expression analysis was carried out using DESeq2 R Bioconductor package [92]. Ranked gene lists were further analyzed by gene set enrichment analysis (GSEA) [93].

### *EGFP* disruption assay for Cas9 activity

Parental cells and Cas9-expressing cells were infected with a lentivirus carrying an *EGFP-2A-puroR* cassette and a sgRNA targeting *EGFP* by spinfection in 12-well plates at 1000 xg for 2 hr, in the presence of 8 µg/µL polybrene. Plates were then returned to 37 °C with 5% CO_2_. Cells were incubated overnight and then selected by puromycin (3 μg/mL, Sigma-Aldrich). Cells were passaged and cultured under puromycin selection for at least 5 days before flow cytometry analysis using an LSRFortessa X20 instrument (BD Biosciences).

### Genome-wide CRISPR synthetic lethality screens

>140 million 786-O-*VHL*^−/−^, 786-O-*VHL*^+^, A-498-*VHL*^−/−^, or A-498-*VHL*^+^ cells were infected with the Avana sgRNA library [62] at a multiplicity of infection (MOI) of 0.3 by spinfection in 12-well plates at 872 xg for 2 hr, in the presence of 8 µg/µL polybrene (Sigma-Aldrich). Plates were then returned to 37 °C with 5% CO_2_. Cells were incubated overnight and then enzymatically detached using trypsin (ThermoFisher Scientifc). Cells for each of the four biological replicates were pooled and seeded into a 5-chamber CellSTACK (Corning) with 800 mL of fresh medium plus 5 µg/mL blasticidin and 3 µg/mL puromycin. After 3-4 days, cells were detached by trypsinization and counted. 40 million cells (~500-fold library coverage) were pelleted for genomic DNA extraction (as early time passages) and another 40 million cells were seeded into a new 5-chamber CellSTACK for passaging. Cell libraries were passaged every 3-4 days until >16 populations doublings, keeping library cell number at 40 million with each passage to maintain adequate sgRNA diversity. At the end of culture period, 40 million cells were pelleted for genomic DNA extraction (as late passages). Genomic DNA was extracted using the DNA Isolation Kit for Cells and Tissues (Roche), according manufacturer’s instruction. The sgRNA sequences were amplified using the primers (listed below) harboring sequencing adaptors and barcodes. In order to achieve >500X coverage over the Avana library (assuming 6.6 μg of genomic DNA for 1 million cells), we performed 30 separate 100 μL PCR reactions with 10 μg genomic DNA in each reaction using ExTaq DNA Polymerase (Clontech) then combined the resultant amplicons. Samples were then purified with SPRIselect beads (Beckman Coulter) according to manufacturer’s instructions. Samples were quantified, mixed and sequenced on a NextSeq 500 (Illumina) by 75-bp single-end sequencing.

#### Forward primer mix

AATGATACGGCGACCACCGAGATCTACACTCTTTCCCTACACGACGCTCTTCCGATCT(0-8 bp variable length sequence)TTGTGGAAAGGACGAAACACCG

#### Reverse primer

CAAGCAGAAGACGGCATACGAGAT(8 bp barcode)GTGACTGGAGTTCAGACGTGTGCTCTTCCGATCTTCTACTATTCTTTCCCC TGCACTGT

### Data processing and sgRNA sequence analysis

After demultiplexing of reads (bcl2fastq, Illumina), quantification of sgRNA across all samples was done with a custom Perl script. Briefly, users either specify the location of the read containing the sgRNA or the flanking primer sequence (if sgRNA position is not constant in all reads). Sequences between the flanking sequences or by location were extracted and compared to a database of sgRNA for each library. Only perfectly matched reads were kept and used in the generation of count matrix. Normalization between all samples was done using the “median ratio method” [94] implemented in the DESeq2 R Bioconductor package. Hit selection for the primary screen was carried out using Student *t* test for the “second best” method and MAGeCK [64].

### Secondary CRISPR screening

The secondary screening procedure was similar to genome-wide CRISPR screens with minor modifications. 20 million cells were used for lentiviral transduction at the MOI of 0.3. At least 6 million cells (500X coverage) were seeded for cell passaging. 10 million cells were collected at early, intermediate and late time points. Genomic DNA was extracted using Quick-gDNA MidiPrep kit (Zymo Research) according to manufacturer’s instructions. 6 million genomic equivalents (~40 µg genomic DNA) were amplified in 4 separate 100 μL PCR reactions with 10 μg genomic DNA in each reaction and then combined for downstream processing.

### Gene ontology and pathway analysis

Gene ontology and pathway analysis was performed with Ingenuity Pathway Analysis (QIAGEN) on the 350 *VHL* synthetic lethal genes. Fisher’s exact test was used to calculate a *P* value determining the probability that each biological function assigned to that data set was due to chance alone.

### Cell proliferation assay for target validation

Cas9-expressing cells were infected with lentiviral constructs carrying *mCherry* and gene-targeting sgRNAs at high MOI by spinfection in 12-well plates at 1000 xg for 2 hr, in the presence of 8 µg/µL polybrene. Plates were then returned to 37 °C with 5% CO_2_. Cells were incubated 48 hr and then trypsinized. Cells were then counted and seeded into 96-well plates. The plates were scanned by Incucyte ZOOM live cell imaging system (Essen Bioscience) every 12 hr. Proliferation of the infected cells was analyzed under red fluorescence automated imaging mode.

### Competitive growth assay

Cas9-expressing cells were transduced with a lentivirus expressing mCherry and a gene-specific sgRNA at high MOI by spinfection in 12-well plates at 1000 xg for 2 hr, in the presence of 8 µg/µL polybrene. Plates were then returned to 37 °C with 5% CO_2_. 48 hr later, cells were trypsinized and seeded into 96-well plates together with the cells expressing vexGFP and a non-targeting control sgRNA. The percentage of mCherry-positive cells were measured at day 2, 6, 10 and 14 post-transduction and normalized to the percentage of mCherry-positive cells at day 2. Flow cytometry analysis was performed on 96-well plates using an LSRFortessa X20. For the trolox rescue experiment, 100 µM trolox (BioVision) were supplemented to the growth medium since day 2 post-transduction.

### Detection of ROS

Cells were plated into white-walled 96-well plates (Corning) at a density of 5,000 cells in 80 μL of culture medium and incubated overnight at 37 °C with 5% CO_2_. Culture medium or ROS inducers such as *tert*-butyl hydroperoxide (Sigma-Aldrich), menadione (Enzo) and pyocyanin (Sigma-Aldrich) were added to the cells in 20 μL solution. 6 hr later, intracellular H_2_O_2_ was detected and quantified using the ROS-Glo H_2_O_2_ assay (Promega) according manufacturer’s instruction.

### Study on the effects of selenium depletion

The isogenic *VHL*^−/−^ and *VHL*^+^ 786-O cells were plated into white-walled 96-well plates at a density of 5,000 cells in 80 μL of ITA-RPMI medium [95], which is made by supplementing RPMI Medium 1640 with 5 µg/mL human insulin (Sigma-Aldrich), 5 µg/mL human transferrin (Sigma-Aldrich), 92 nM FeCl_3_ (Sigma-Aldrich), and 2.5 mg/mL bovine serum albumin (ThermoFisher Scientific). ROS signals were measured after 48 hour incubation at 37 °C with 5% CO_2_. As control groups, cells were grown in ITA-RPMI medium plus 6.25 nM sodium selenite (Sigma-Aldrich) or RMPI Medium 1640 plus 10% FBS for 48 hours before ROS detection.

### Cell viability assay for drug treatment

Cells were plated into black-walled 384-well plates (Corning) together with mimethyl sulfoxide or chemicals at various concentrations (3-fold dilution). Cell viability was quantified using Cell Titer-Glo Luminescent Cell Viability Assay (Promega) according manufacturer’s instruction.

### Primary tumor sample analysis and survival analysis

We used TCGA data sets of gene expression data (Illumina Hiseq RNA sequencing data) for kidney renal clear cell carcinoma samples and patients’ clinical record (http://gdac.broadinstitute.org/). Differential Expression analysis was carried out using DESeq2 R Bioconductor package [92] in order to identify the genes that are overexpressed in primary tumors versus normal tissues, using a cutoff of *P* value < 0.05. For the survival analysis, we use ‘time-to-event’ univariate Cox proportional hazard model on each gene to determine the impact. Kaplan-Meier plots also show the comparisons between curves by log rank test. General survival analysis over the gene list utilized median *z* score of gene expressions among “Tumor” and “Normal” groups.

## Acknowledgements

We thank David Root, Olivia Bare, John Doench, Xiaoping Yang, Thomas Green, and other members of the Genetic Perturbation Platform of The Broad Institute (Cambridge, MA) for sgRNA library construction and other reagents.

## Author contributions

N.S. designed research, generated and characterized ccRCC isogenic cell line pairs, conducted RNA-seq, primary and validation CRISPR screening, performed target validation and mechanistic studies, analyzed data, and wrote the manuscript. S.P. assisted with primary CRISPR screening. C.L. assisted with data analysis of RNA-seq and CRISPR screening. J.E.H assisted with designing research and assay development. M.H. conducted sgRNA deconvolution. M.H. assisted with data analysis of RNA-seq and TCGA clinical data. M.H.D. assisted with RNA-seq and sgRNA deconvolution. D.M. tested cellular response to small molecule DDR inhibitors. S.N.A. assisted with assay development. C.P.M assisted with designing research. N.C. designed and supervised the research and wrote the manuscript.

## Supplementary information

### Supplementary figures

Supplemental Fig. S1. Representative flow cytometry histograms of the *EGFP*-disruption assay.

Supplemental Fig. S2. Gene set enrichmnt analysis of differentially expressed genes in *VHL*^−/−^ versus *VHL*^+^ ccRCC cells.

Supplemental Fig. S3. *VHL* status does not affect cell proliferation in standard cell culture conditions.

Supplemental Fig. S4. Heatmaps of Pearson’s correlations of the library distribution from two pairs of ccRCC isogenic cell lines.

Supplemental Fig. S5. Pearson’s correlations between biological replicates in two pairs of ccRCC cell lines at different time points.

Supplemental Fig. S6. Density plots of log2-transformed sgRNA counts in different samples. All sgRNAs are shown in curves filled in grey. sgRNAs targeting essential genes are shown in red curves. sgRNAs targeting nonessential genes are shown in blue curves. Non-targeting control sgRNAs are shown in green curves.

Supplemental Fig. S7. Inactivation of *SEPHS2* or *PSTK* using two individual sgRNAs led to selective growth inhibition in *VHL*^−/−^ cells. Data represent mean ± SD (*n* = 3). *P* values were derived from *t* tests on area under the curve.

Supplemental Fig. S8. Representatives of the flow cytometry plots of the fluorescence competitive growth assay in A-498 cells. The isogenic *VHL*^−/−^ and *VHL*^+^ cells were cultured in the standard growth medium with or without 100 µM trolox.

Supplemental Fig. S9. Representatives of the flow cytometry plots of the fluorescence competitive growth assay in 786-O cells. The isogenic *VHL*^−/−^ and *VHL*^+^ cells were cultured in the standard growth medium with or without 100 µM trolox.

Supplemental Fig. S10. Inactivation of *CDK1, CHEK1* or *TOP2A* using two individual sgRNAs led to selective growth inhibition in *VHL*^−/−^ cells. Data represent mean ± SD (*n* = 3). *P* values were derived from *t* tests on area under the curve.

Supplemental Fig. S11. Gene set enrichment analysis of differentially expressed *VHL-*SL genes associated with DDR in *VHL*^−/−^ versus *VHL*^+^ ccRCC cells.

### Supplementary tables

Supplemental Table S1. Deseq analysis of the RNA-seq data from A-498-*VHL*^−/−^ and A-498-*VHL*^+^.

Supplemental Table S2. Deseq analysis of the RNA-seq data from 786-O-*VHL*^−/−^ and 786-O-*VHL*^+^.

Supplemental Table S3. MAGeCK analysis of primary CRISPR synthetic lethality screening.

Supplemental Table S4. Differential sgRNA depletion analysis of primary CRISPR synthetic lethality screening.

Supplemental Table S5. *VHL* SL gene lists isolated from two isogenic cell line pairs using two different hit selection strategies.

Supplemental Table S6. Synthetic lethality scores of primary and secondary CRISPR screening.

Supplemental Table S7. MAGeCK analysis of seconday CRISPR synthetic lethality screening.

Supplemental Table S8. FPKM information in A-498 and 786-O isogenic cell pairs.

Supplemental Table S9. Gene lists for Figure 2C.

Supplemental Table S10. Enriched canonical pathways by Ingenuity Pathway Analysis from the 350 *VHL* SL genes.

Supplemental Table S11. IC_50_ values of small molecular inhibitors targeting AURKA, AURKB, ATR, CDK1 and TOP2A in ccRCC isogenic cell pairs.

Supplemental Table S12. 52 *VHL* SL hits associated with DDR are significantly upregulated in primary tumors versus normal solid tissues (TCGA-KIRC). *P* values were derived from *t* test.

