## Supplementary figures for "*VHL* Synthetic Lethality Signatures Uncovered by Genotype-specific CRISPR-Cas9 Screens"

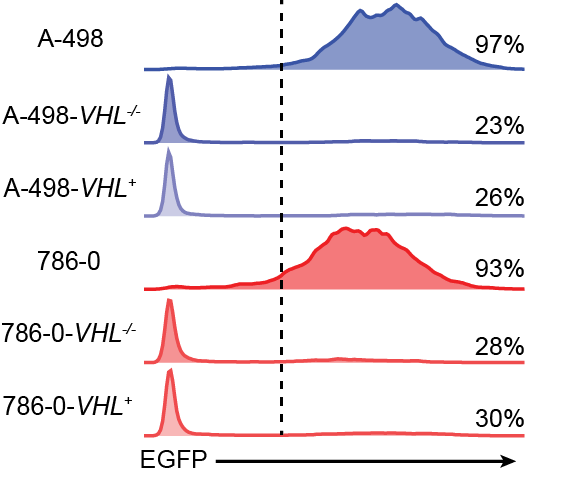

Supplemental Fig. S1. Representative flow cytometry histograms of the *EGFP*-disruption assay.

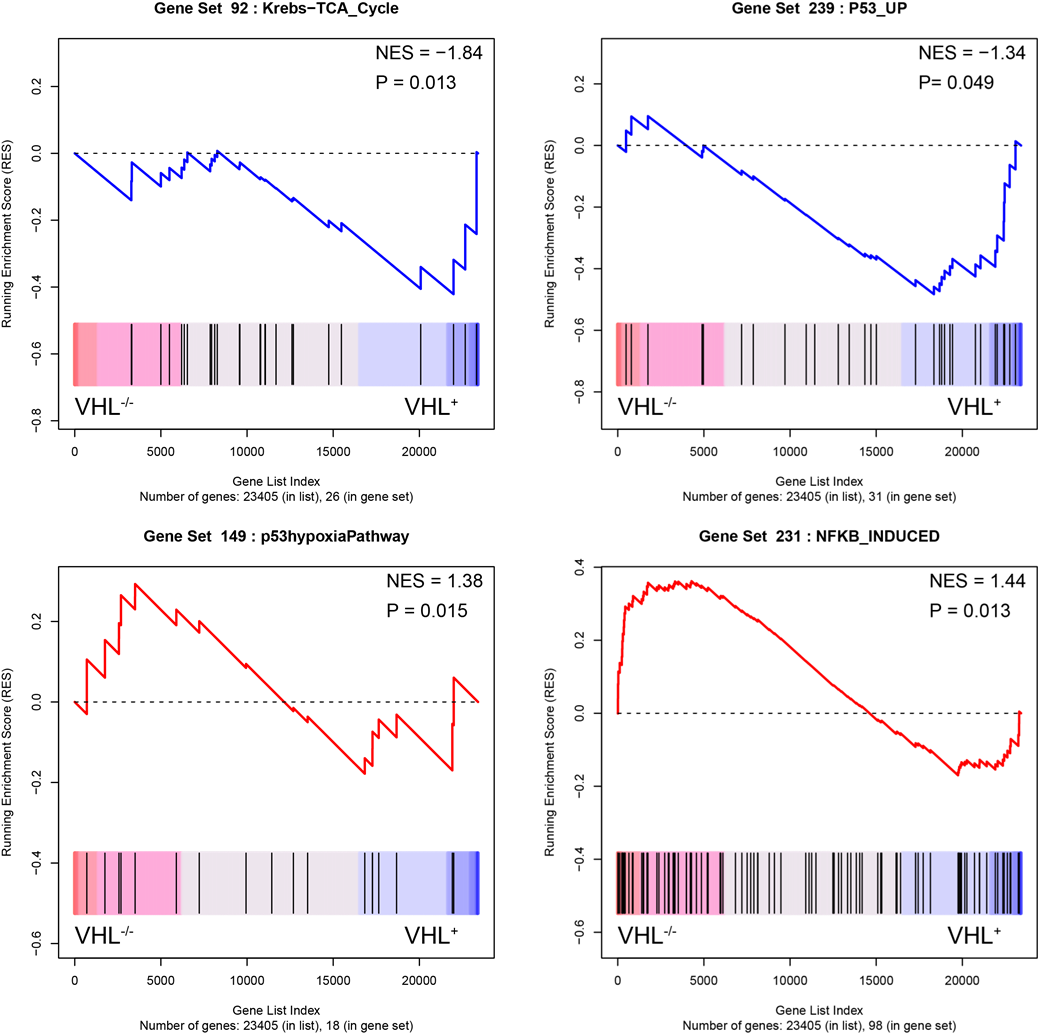

Supplemental Fig. S2. Gene set enrichment analysis of differentially expressed genes in *VHL^-/-^* versus *VHL^+^* ccRCC cells.

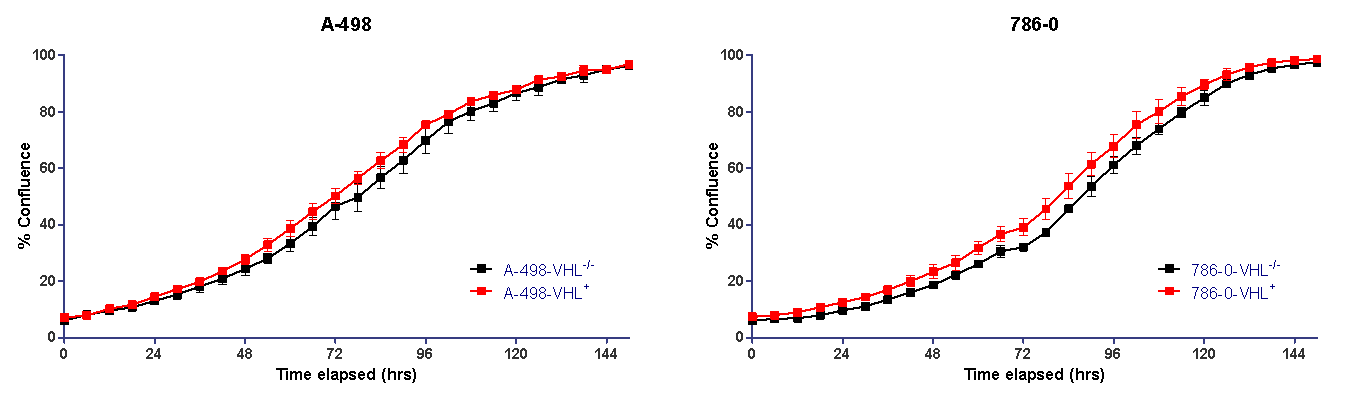

Supplemental Fig. S3. *VHL* status does not affect cell proliferation in standard cell culture conditions.

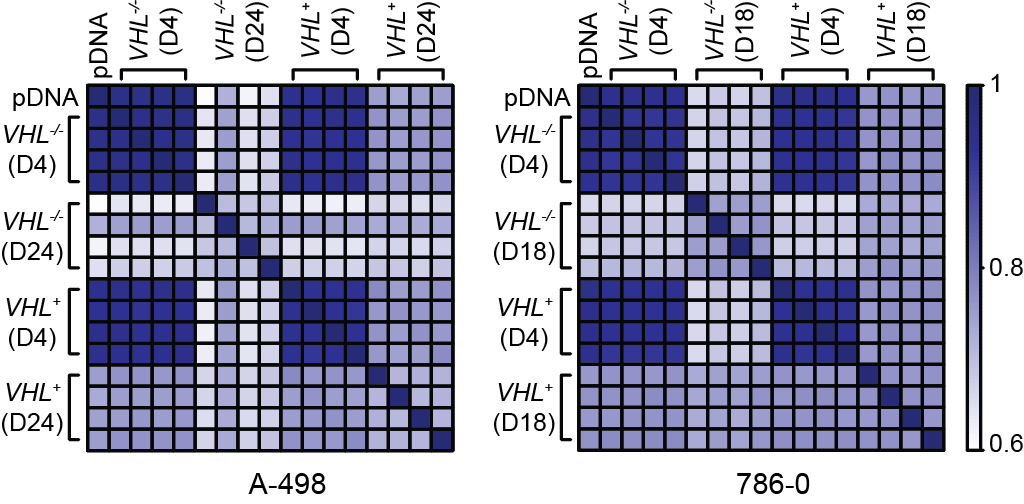

Supplemental Fig. S4. Heatmaps of Pearson’s correlations of the library distribution from two pairs of ccRCC isogenic cell lines.

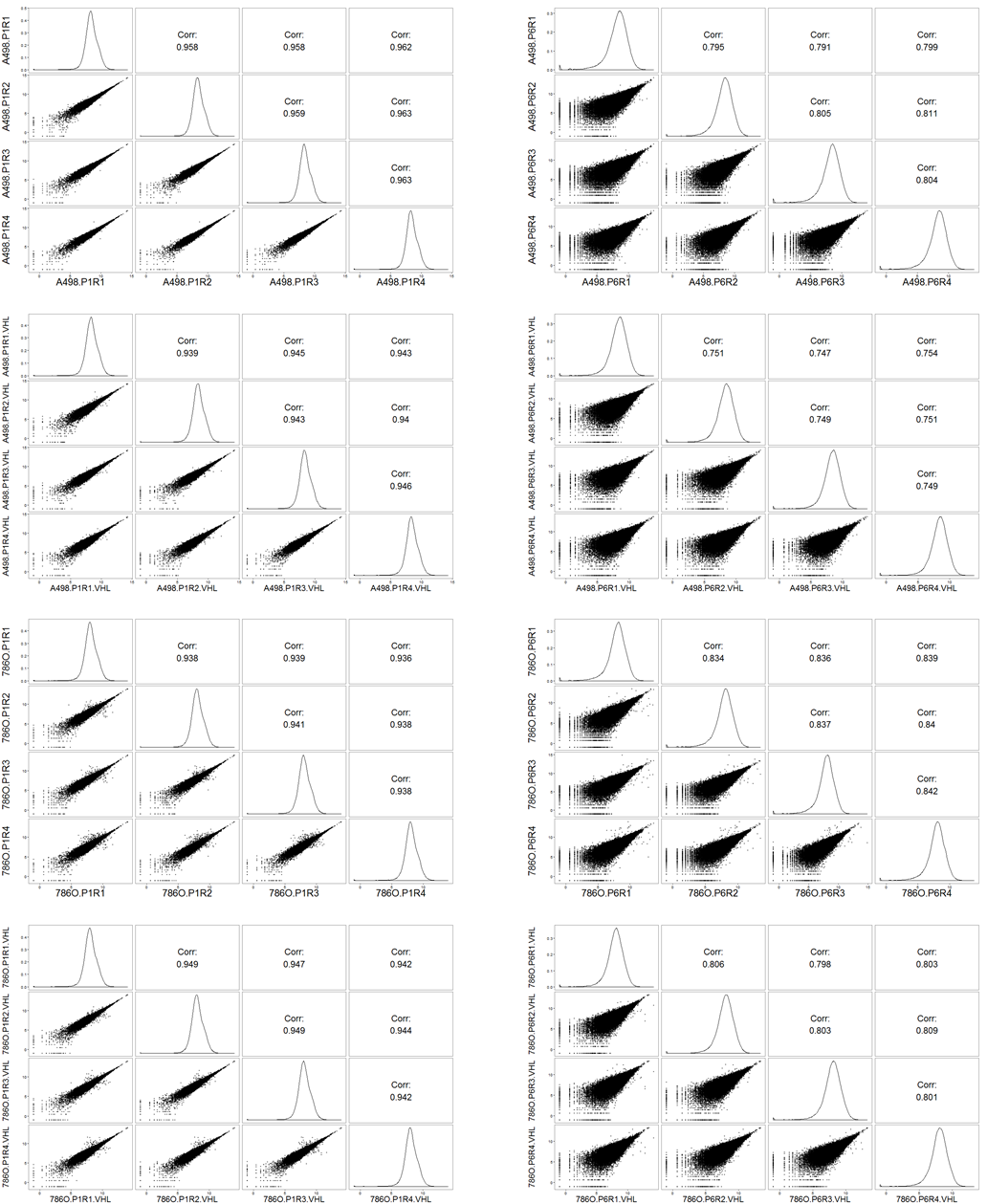

Supplemental Fig. S5. Pearson’s correlations between biological replicates in two pairs of ccRCC cell lines at different time points.

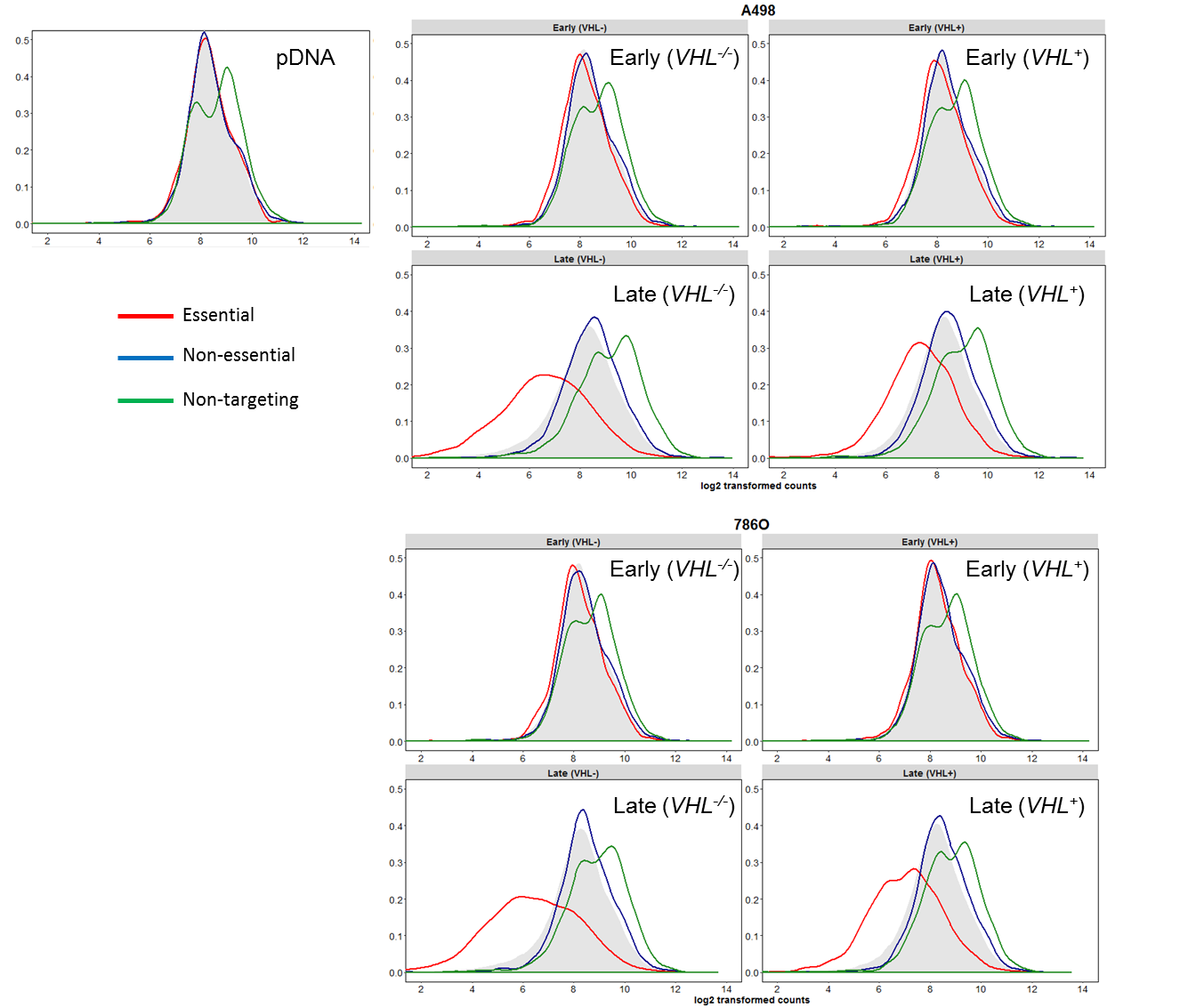

Supplemental Fig. S6. Density plots of log2-transformed sgRNA counts in different samples. All sgRNAs are shown in curves filled in grey. sgRNAs targeting essential genes are shown in red curves. sgRNAs targeting nonessential genes are shown in blue curves. Non-targeting control sgRNAs are shown in green curves.

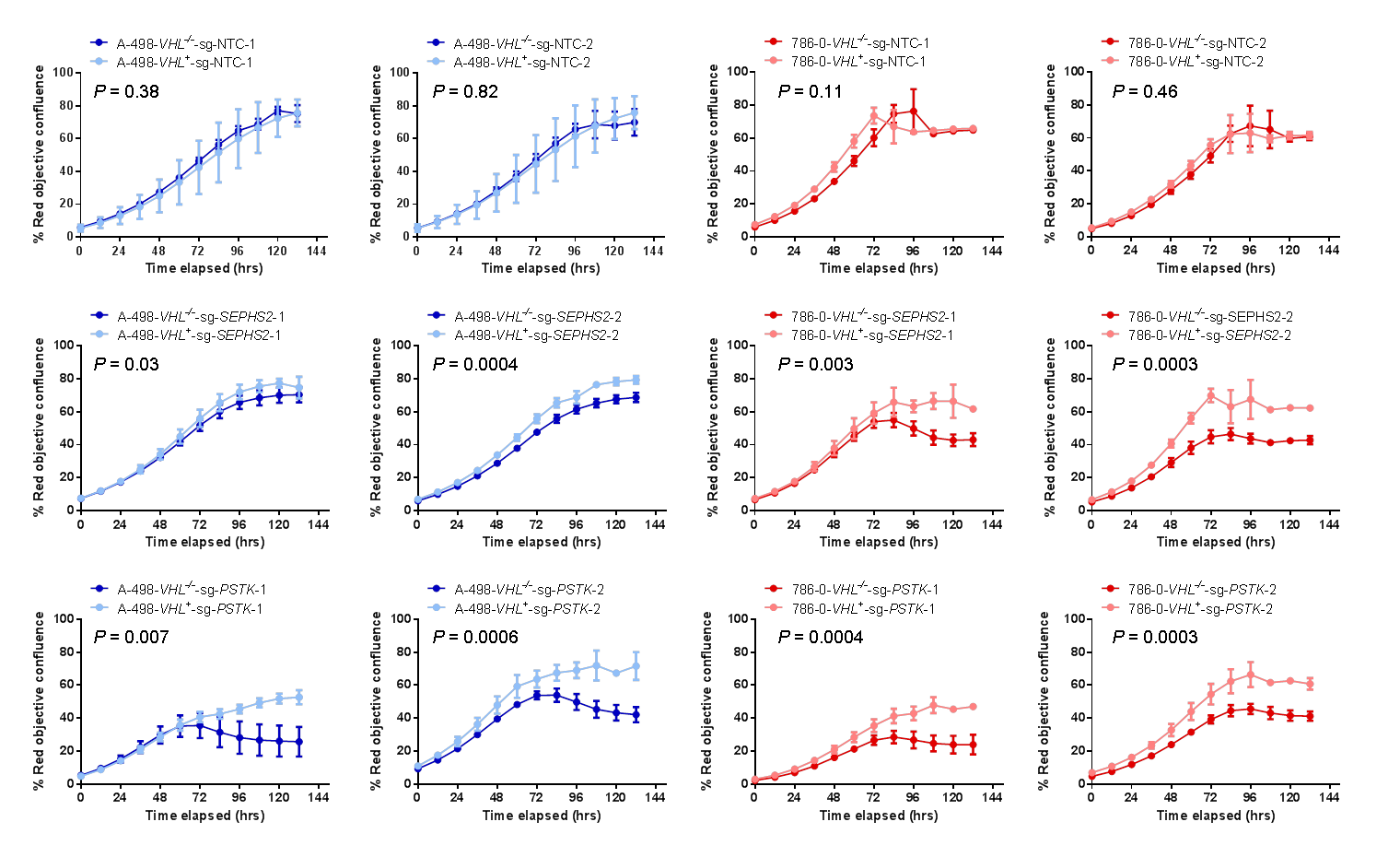

Supplemental Fig. S7. Inactivation of *SEPHS2* or *PSTK* using two individual sgRNAs led to selective growth inhibition in *VHL^-/-^* cells. Data represent mean ± SD (*n* = 3). *P* values were derived from *t* tests on area under the curve.

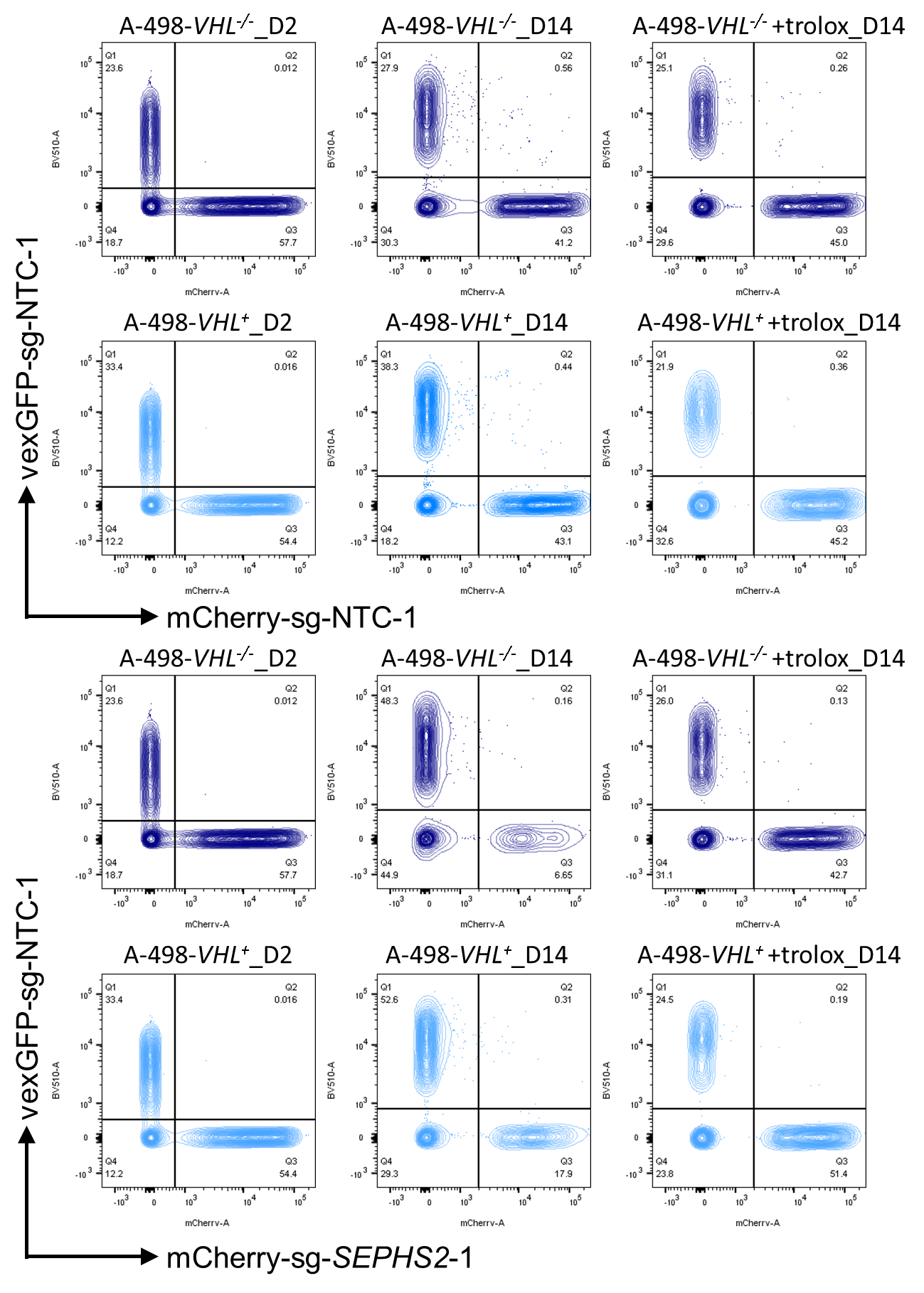

Supplemental Fig. S8. Representatives of the flow cytometry plots of the fluorescence competitive growth assay in A-498 cells. The isogenic *VHL^-/-^* and *VHL^+^* cells were cultured in the standard growth medium with or without 100 µM trolox.

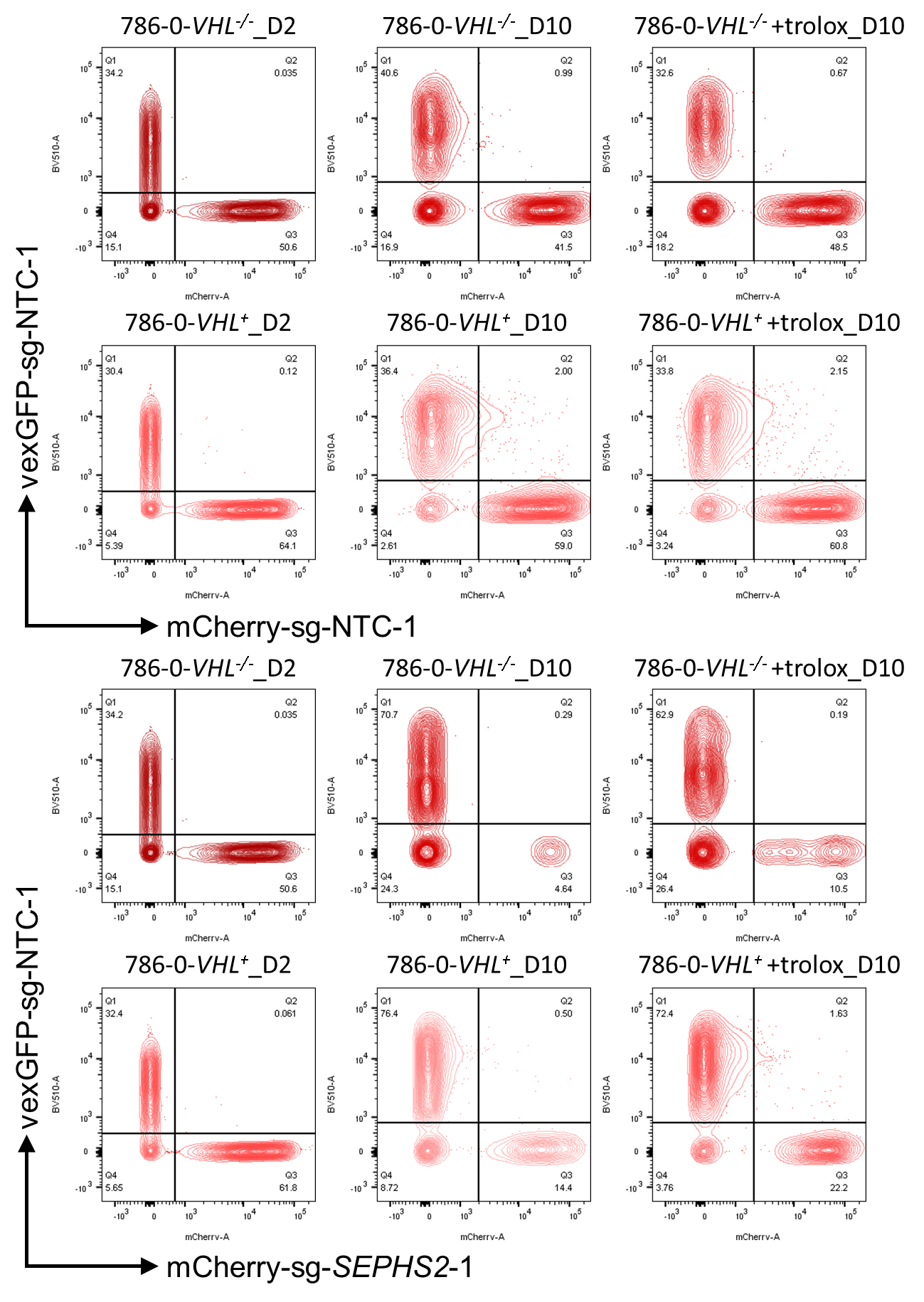

Supplemental Fig. S9. Representatives of the flow cytometry plots of the fluorescence competitive growth assay in 786-O cells. The isogenic *VHL^-/-^* and *VHL^+^* cells were cultured in the standard growth medium with or without 100 µM trolox.

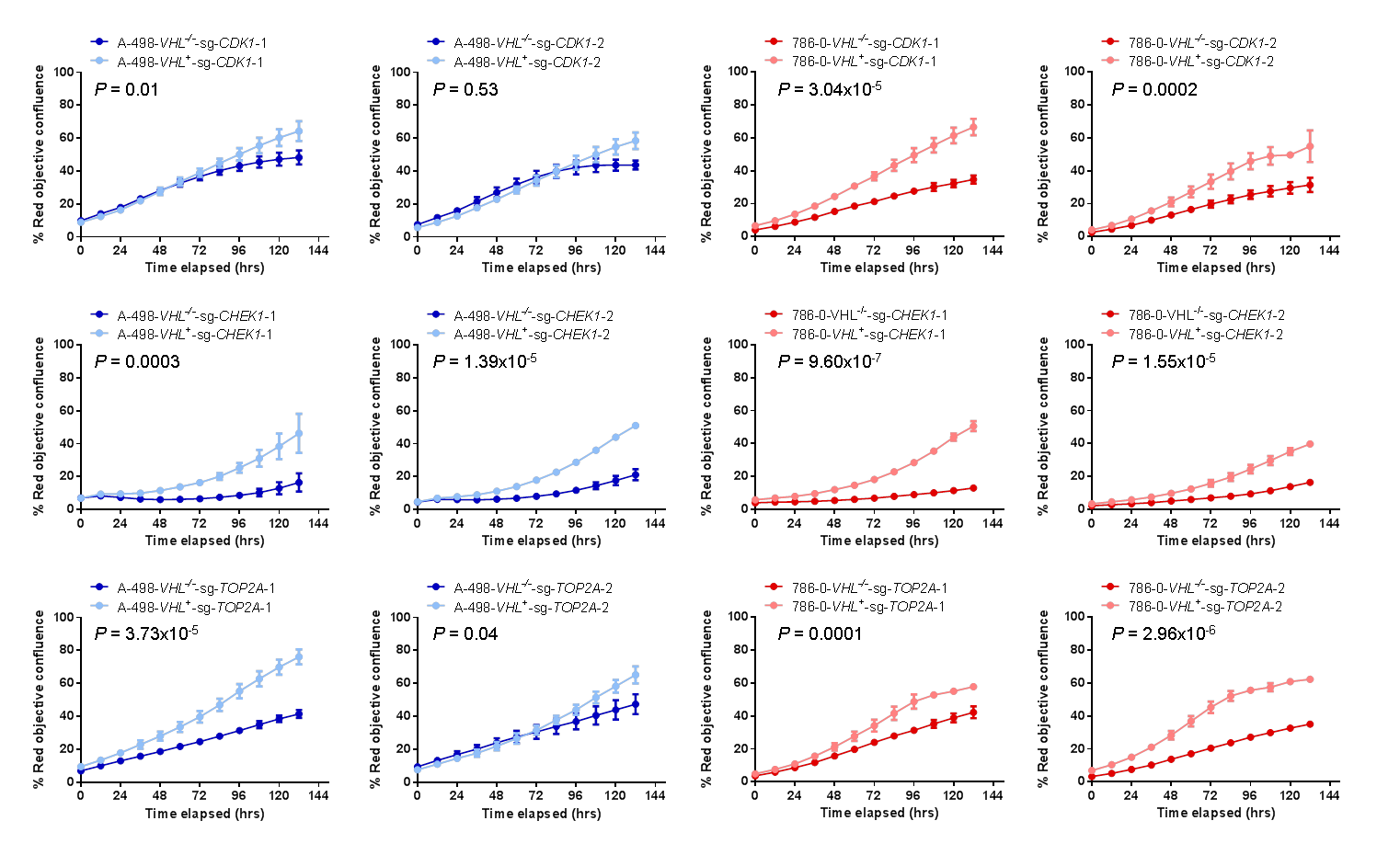

Supplemental Fig. S10. Inactivation of *CDK1, CHEK1* or *TOP2A*  using two individual sgRNAs led to selective growth inhibition in *VHL^-/-^* cells. Data represent mean ± SD (*n* = 3). *P* values were derived from *t* tests on area under the curve.

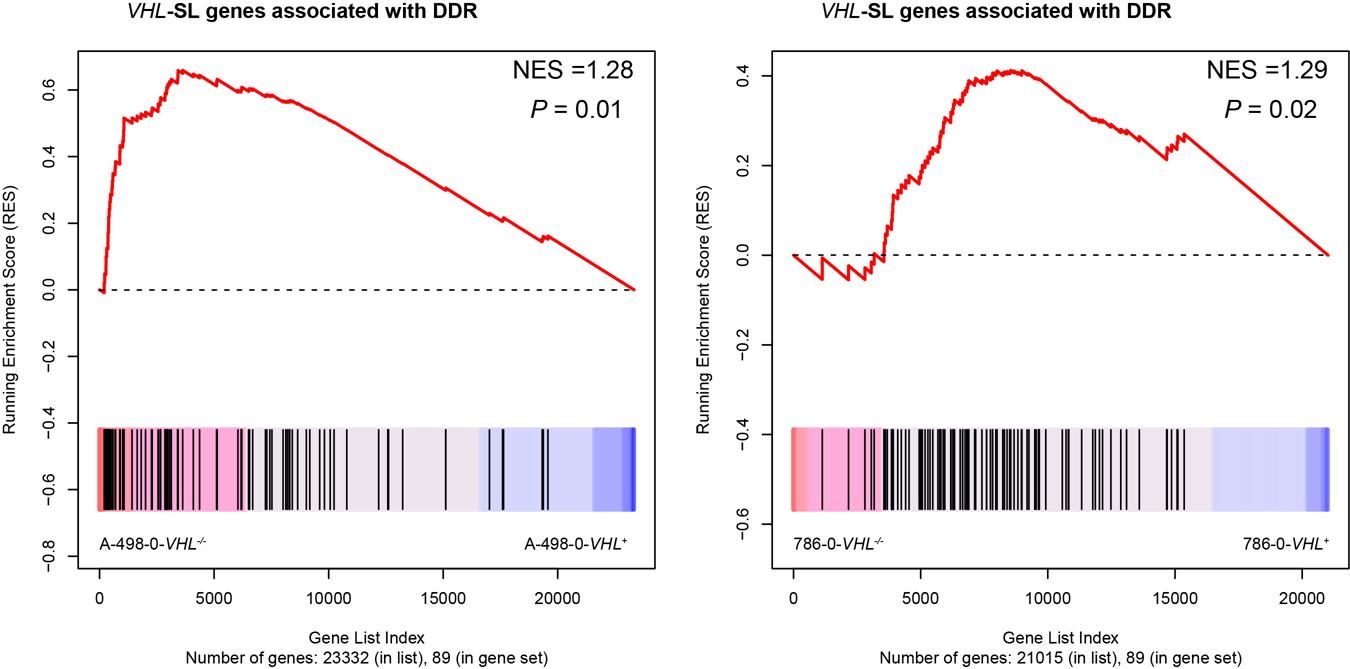

Supplemental Fig. S11. Gene set enrichment analysis of differentially expressed *VHL-*SL genes associated with DDR in *VHL^-/-^* versus *VHL^+^* ccRCC cells.
